# A deep learning approach for rational affinity maturation of anti-VEGF nanobodies

**DOI:** 10.1101/2025.10.20.683442

**Authors:** Gaëlle Verdon, Laurent David, Alexandre de Brevern, Yasser Mohseni Behbahani

## Abstract

Nanobodies offer several advantages over conventional antibodies due to their lower immunogenicity, enhanced stability, and superior tissue penetration, making them promising candidates for cancer therapy. In this study, we employ deep learning algorithms to design anti-VEGF nanobodies via affinity maturation. Our approach integrates structure-guided mutational modeling and systematic measurement of binding affinity and stability for rational optimization of Complementarity Determining Regions. In addition, we developed a sequence-based melting temperature predictor for nanobodies, ensuring stability of the designed mutants. Our method achieves energy reductions up to -4.92 kcal/mol. Our melting temperature predictor demonstrated a Pearson correlation coefficient of 0.772. These findings emphasize the potential of computational approaches for nanobody affinity maturation and stability prediction, paving the way for more effective therapeutic designs.

## 1 Introduction

Nanobody or VHH (Fig. **1**) is the single variable domain of heavy-chain-only antibodies (HCAbs) which retains full binding capacity despite its compact size (15kDa, 10 times smaller than antibodies) [2]. Due to their small size, VHHs have superior stability, enhanced tissue permeability and rapid blood clearance compared to conventional antibodies [29, 17], making them particularly effective for targeting dense tumors and crossing the blood-brain barrier [29]. They rely solely on three complementarity-determining regions (CDRs), including an extended CDR3 loop (16–24 amino acids vs. ∼ 10 in antibodies) [9], to recognize antigens with high specificity and affinity. This elongated CDR3 allows VHHs to access sterically restricted or cryptic epitopes that are often inaccessible to bulkier antibodies, such as enzyme active sites or viral canyons [11, 29, 9, 26]. These favorable biophysical and pharmacokinetic properties make VHHs ideal candidates for anti-angiogenic therapeutic solutions [5, 13].

**Figure 1:**
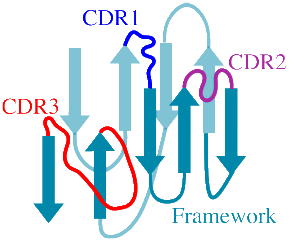
VHH secondary structure.

Vascular endothelial growth factor (VEGF) regulates angiogenesis and is often overexpressed in cancer, promoting tumor growth [25]. Blocking the interaction between VEGF and its receptor (VEGFR) is a key therapeutic approach in oncology [5]. A promising strategy involves designing protein-based binders, such as VHHs, to disrupt this interaction. Recent advancements in computational biology have transformed protein engineering, with deep learning approaches emerging as a powerful alternative to traditional experimental methods. These approaches accelerate affinity maturation between the binder and the target and improve efficiency by exploring the mutational landscape to identify beneficial variations [14, 10]. They contribute to multiple stages of affinity maturation, including binder design, structural modeling, and assessment of mutation-induced changes in binding affinity 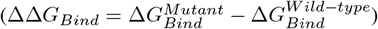 between the wild type and the mutant complexes.

39th Conference on Neural Information Processing Systems - Machine Learning in Structural Biology Workshop A class of deep learning-based methods is protein language models (pLM), which extract biophysical and evolutionary information that determine protein structure and function [7, 21, 15]. Sequence-based deep learning approaches like MuLAN rely on pLM and offer a fast and accurate prediction of ΔΔ*G*_*Bind*_ in protein-protein interactions [16]. However, CDR sequences are highly diverse and are not concerned by the notion of conservation during evolution. This limits the ability of pLM to extract sufficient predictive information from interfaces where these regions are involved. Another class operates on protein design, addressing the inverse folding problem: identifying amino acid sequences that fold into a given 3D backbone structure. ProteinMPNN, a structure-based graph neural network, is an example of these methods applied to the problem of *de novo* binder design [3]. AbMPNN is a fine-tuned version of ProteinMPNN trained specifically for antibodies; it focuses on determining and optimizing the CDR3 [4]. For protein structure prediction, AlphaFold-Multimer [8] and AlphaFold3 [1] accurately reconstruct protein complexes and provide insights into binding mechanisms. ColabFold is a faster implementation of AlphaFold-Multimer, which optimizes inference and accessibility while maintaining similar performance [18]. Structure-based tools like DLA-Mutation [19] and DDgPredictor [24] accurately estimate ΔΔ*G*_*Bind*_, with the latter being especially effective for antibody-antigen interactions.

Thermostability is essential for VHH functionality and design feasibility. It is shown that the VHH’s CDR loops play a crucial role in its stability [6]. TemBERTure, a deep learning model, predicts protein melting temperature (*T*_*m*_) by leveraging pLM protBERT-BFD [7], fine-tuned through an adapter-based approach [22]. It shows great promise in the *T*_*m*_ prediction of a wide range of proteins.

This study aims to develop a deep learning-based affinity maturation framework for the rational design of anti-VEGF nanobodies with improved binding affinity and thermal stability. ColabFold structural predictions enable us to evaluate the effect of sequence variants on the VHH-VEGF complex, and DDgPredictor provides precise estimates of ΔΔ*G*_*Bind*_. To ensure the biophysical robustness of VHH variants, we trained a VHH-specific model derived from TemBERTure to predict the thermostability of new VHH designs. Our integrative approach enables us to explore the mutational landscape in the CDR3 loop and beyond at high throughput, balancing enhanced binding with stability preservation.

## 2 Materials and Methods

### 2.1 Affinity maturation pipelines

After selecting antibodies and VHH that bind VEGF, their CDRs are assigned using human expertise. This step is followed by introducing mutations on the CDR3 using ProteinMPNN and exhaustive mutational scanning. ProteinMPNN generates CDR3 sequences conditioned to the VHH-VEGF complex structure. The Structure of wild-type and mutant are predicted using ColabFold, which produces multiple models per sequence; due to computational limits, only the best structure is kept when the sequence does not change. The ΔΔ*G*_Bind_ is then predicted using DDgPredictor to identify favorable mutants (ΔΔ*G*_*Bind*_ *<* 0). We developed three pipelines for affinity maturation.

#### Wild-type based maturation

Starting from the VHH-VEGF complex structure with identified CDR regions, we generated 10,000 variants by introducing multiple point mutations within the CDR3 using ProteinMPNN. A redundancy reduction step of the resulting mutant sequences is performed, keeping them unique. We also conducted exhaustive single-point mutations at every position of the CDR3 without considering ProteinMPNN (**Fig. S1**).

#### Structure-adapted maturation

We adapt the wild-type based maturation pipeline to account for mutation-induced conformational variability. At each position of CDR3, we generate a single-point mutation following the ProteinMPNN probability output. We predict the structure and ΔΔ*G*_*Bind*_ following each positional mutation (**Fig. 2**). This allows us to adjust the structure along the design. However, since mutations are performed sequentially, one after the other, scanning the CDR3, the generation of mutations on the final amino acid is influenced by the preceding ones. As a result, not all combinations of mutations may be explored. To address this, we add an option to continue the process iteratively. Each iteration begins with the structure and sequence produced at the final step of the previous iteration (*i*.*e*. the last position of the CDR3 sequence). This iterative process explores more thoroughly the mutational landscape and eventually discovers more favorable sequences. We add another option to guide the mutations towards higher complex stability (guided maturation process). We retain mutations that lower ΔΔ*G*_*Bind*_ before moving to the next position (**Fig. 2**).

**Figure 2:**
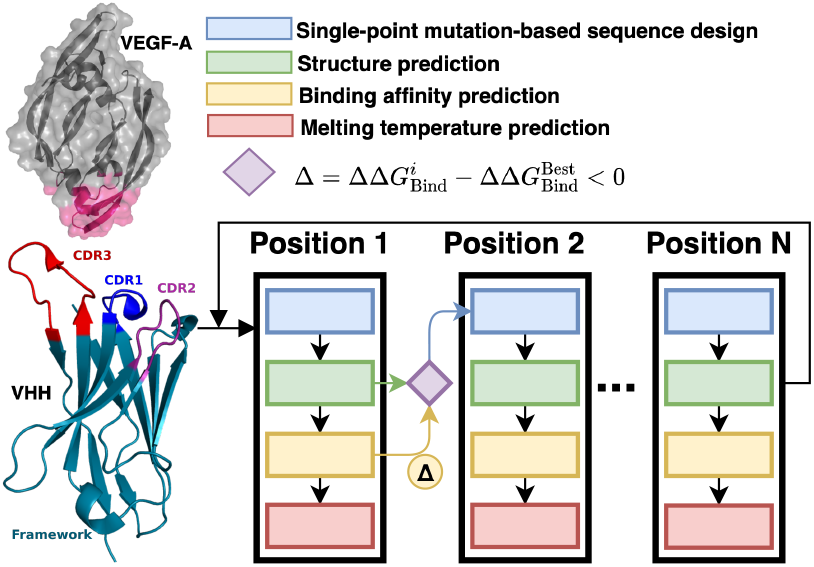
Structure-adapted maturation pipeline. A mutation is applied to VHH at position i of CDR3 following the amino acid substitution generated by ProteinMPNN. After each mutation, the pipeline determines the complex structure and predicts ΔΔ*G*_*Bind*_. The subsequent amino acid substitution at position i+1 is determined by the mutant structure generated for position i. One can repeat the process several iterations. In each iteration, process k+1 starts with the structure and sequence of the mutant complex designed for position N from process k. In the guided maturation pipeline, the structure and the sequence of the mutant are kept only if the process finds a more favorable sequence 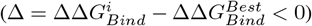

### 2.2 Thermostability prediction

We finetuned the TemBERTure best model (model replica 2) for VHH’s *T*_*m*_ prediction using the NbThermo dataset. The learning rate was optimized while the other parameters were kept at their default values (dropout in the adapter head is set to 0.2, no weight decay, and warmup ratio of 0). We also finetuned the protBERT model following the adapter architecture proposed by the TemBERTure. The learning rate, the number of layers, the dropout, and the activation function of the adapter head were optimized, while a warmup ratio of 0 and no weight decay were kept as default.

### 2.3 Data

#### Affinity maturation

The two nanobodies and the antibody used in this study come from SabDab-nano and SabDab, respectively, two publicly available databases containing experimentally determined antibody and nanobody structures [23]. The VHH-VEGF complexes 3P9W and 5FV2 were extracted using “VHH” and “vascular endothelial growth factor” as *Antibody type* and *keyword query*, respectively. The Antibody-VEGF complex 6BFT was extracted using “Fv” and “vascular endothelial growth factor A” as *Antibody type* and *keyword query*, respectively. The 5FV2 complex consists of a VHH and a dimeric VEGF, while the 3P9W complex contains a VHH in interaction with a monomeric VEGF.

#### Thermostability prediction

For training our model, we used the NbThermo dataset, containing *T*_*m*_, amino acid sequences, and annotated CDRs of 564 VHHs [27]. We kept 513 non-redundant VHH sequences with available *T*_*m*_. To prevent data leakage, we ensured that there was no redundancy of CDRs across our training, validation, and test sets. This resulted in the test, validation, and training sets of 52, 51, and 410 samples, respectively.

## 3 Experiments and Results

We applied the wild-type based approach on complexes 3P9W, 5FV2, and 6BFT and generated 10’000 CDR3 sequences (multiple point mutations) for each, among which 197, 185, and 5811 were unique for 3P9W, 5FV2, and 6BFT, respectively. We further filtered the mutations on 6BFT to include sequences generated more than 6 times by ProteinMPNN to reduce computational costs. This filtering further reduced the number of mutants to 229. The greater diversity of mutations observed for the 6BFT antibody complex may result from an increased number of possible mutation combinations, facilitated by the presence of both heavy and light chain CDR3, rather than simply one CDR3. Within the unique mutations generated, 68, 57, and 142 mutations were favorable for 3P9W, 5FV2, and 6BFT complexes, respectively.

We ran the structure-adapted maturation process on the CDR3 of 5FV2 (positions 97 to 109 of VHH) ten independent times (**Fig. 3A** and **S6**). For ease of visualization, we selected 6 processes out of 10, from which 5 converged to favorable mutants (**Fig. 3A**). For all generated mutant sequences, the corresponding melting temperature is always lower than the wildtype (71.5 °C).

**Figure 3:**
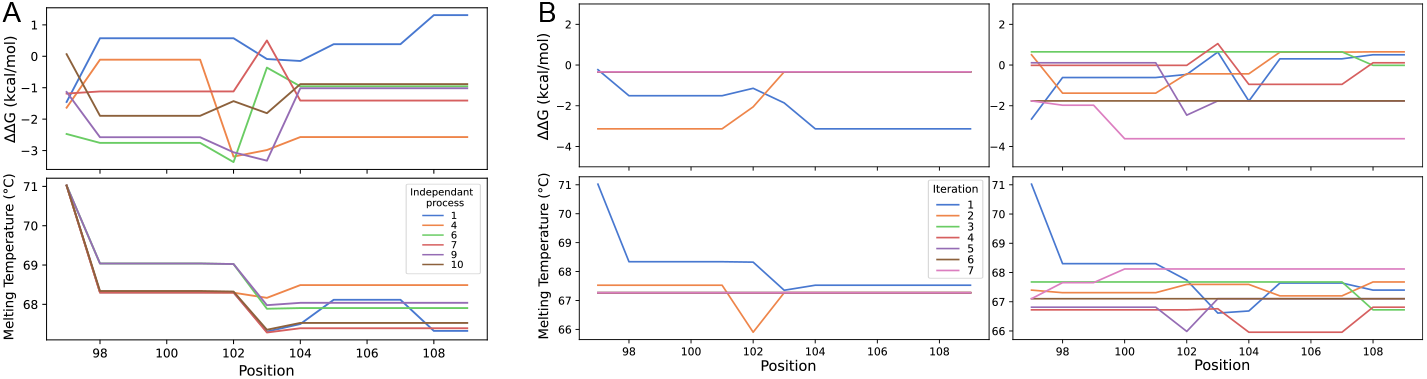
Evolution of the predicted ΔΔ*G*_*Bind*_ (top row) and *T*_*m*_ (bottom row) through mutant positions within the CDR3 of the complex 5FV2. **(A)** Each curve corresponds to a selected independent design process (see **Fig. S6** for all results). **(B)** Evolution over 7 iterations for two independent iterative processes (left and right). On the left, while iteration 1 (blue) reduces ΔΔ*G*_*Bind*_, iteration 2 (orange) increases it back to close to 0 (almost no improvement), and subsequently, the sequence and structure remain the same for all following iterations. On the right, ΔΔ*G*_*Bind*_ fluctuates around -1 *kcal/mol*, throughout iterations, and at the end of the last iteration, we reach the most negative ΔΔ*G*_*Bind*_ value, around -4 *kcal/mol*. The X-axis corresponds to the CDR3 positions.

Similarly, we ran the iterative process on the CDR3 of complex 5FV2 ten independent times. Each process has 7 iterations. Almost all resulting sequences are favorable (**Fig. 3B** and **S7**). We also observed that the final mutant obtained may not necessarily be the most favorable during the iterative process. This could be due to divergences between mutations judged as favorable by ProteinMPNN and the ones predicted as favorable by DDgPredictor. We observed lower *T*_*m*_ for mutants compared to the wild-type. All along the iterative process, *T*_*m*_ fluctuates within 3°C.

We explored the space of single-point mutations for the two VHH-VEGF complexes. We observed that certain wild-type positions support multiple favorable amino acid substitutions (**Fig. 4A**). To assess whether ProteinMPNN performance aligns with the predicted mutational landscape associated with exhaustive single-point mutations, we ran ProteinMPNN on each position 10 independent times and averaged the output probability vectors. There are differences in the choice of amino acid substitutions between ProteinMPNN and the ΔΔ*G*_Bind_ predictions from exhaustive single-mutational scanning (**Fig. 4A**). The charge distributions on the interface of wild-type and mutant VHH-VEGF structures align with the DDgPredictor predictions. Building on this observation, we decided to run the guided maturation process on the CDR3 of 5FV2 ten independent times, by evaluating the predicted ΔΔ*G*_*Bind*_ after each mutation to converge to the highest complex stability. We observed a progressive decrease of ΔΔ*G*_Bind_, reaching -4.92 *kcal/mol* at the end of affinity maturation for complex 5FV2 (**Fig. 4B**). The most favorable mutants, reaching ΔΔ*G*_*Bind*_ values smaller than -4 kcal/mol (processes 2, 3, 5, 9, 10), also reach higher *T*_*m*_. The least favorable mutant (process 4) reaches the lowest *T*_*m*_ while the most favorable one (process 3) reaches the highest one.

**Figure 4:**
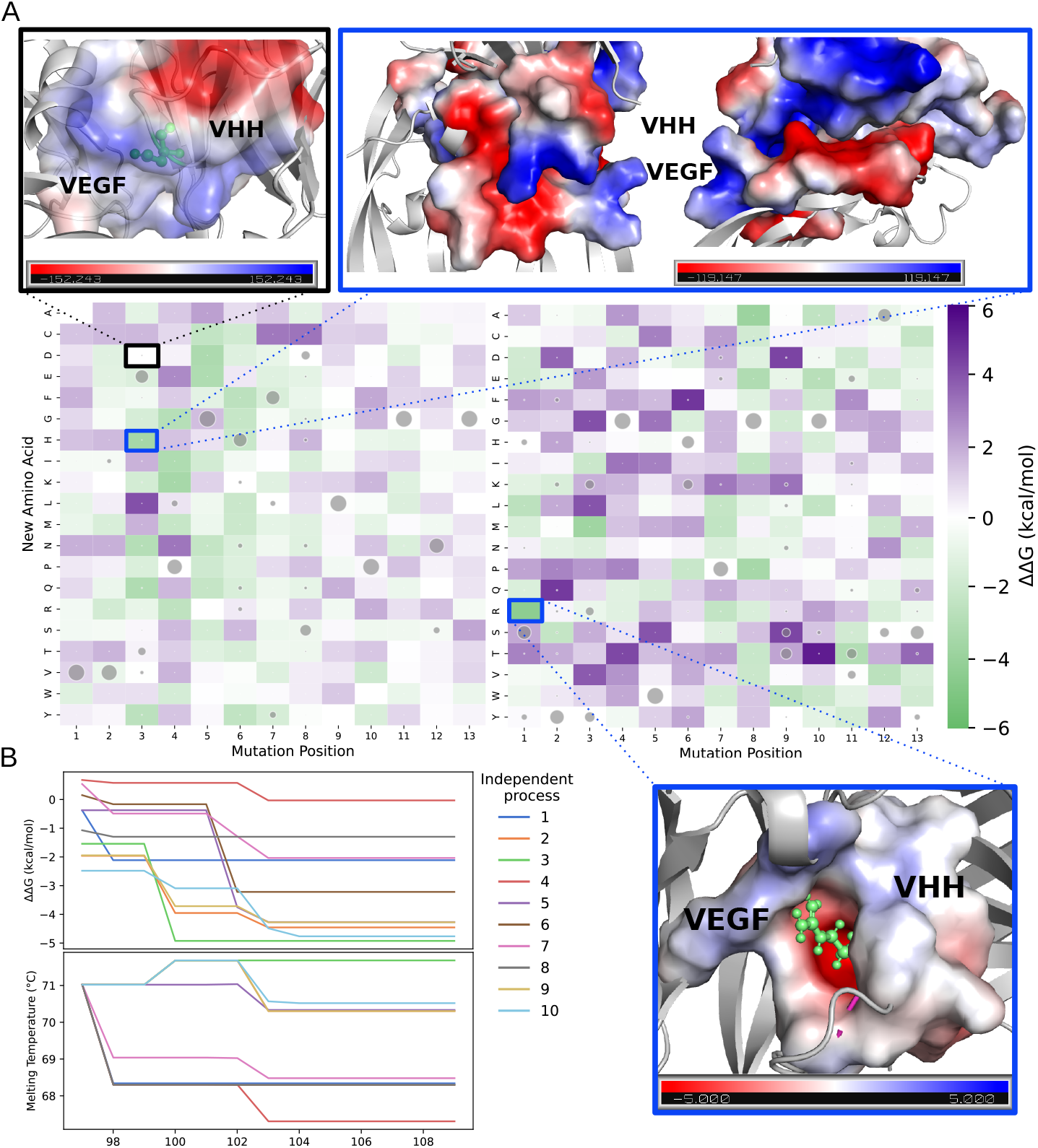
**(A)** Predicted ΔΔ*G*_*Bind*_ for exhaustive single point mutation on the CDR3 of 5FV2 (left) and 3P9W (right). Gray circles represent the mean probability of substitution, generated at Softmax temperature 0.25 for 5FV2 and 0.4 for 3P9W, obtained over 10 runs by ProteinMPNN for each position. In the structure of complexes, the mutant amino acid is shown in green, and the electrostatic charge distribution goes from negative charge (red) to positive charge (blue). At the interface of 5FV2, the surface of VEGF exhibits a predominantly negative electrostatic potential, which creates an unfavorable environment for the aspartic acid (D) residue at position 3 of CDR3. Substituting this residue with histidine (H) enhances binding affinity by introducing a more favorable electrostatic interaction. **(B)** Evolution of the predicted ΔΔ*G*_*Bind*_ (top) and *T*_*m*_ (bottom) across CDR3 mutations of VHH in 5FV2 during guided maturation over 10 independent processes. All processes reach lower ΔΔ*G*_*Bind*_ values, and process 3 gives the lowest value. All *T*_*m*_ predictions are lower compared to wild-type at the end of each process, except for process 3. The X-axis corresponds to the CDR3 positions.

Since ProteinMPNN and DDgPredictor perform differently in identifying favorable mutations, we sought to determine whether AbMPNN can align with DDgPredictor and generate diverse mutations. We calculated the mean of probability vectors of amino acid substitutions (computed at Softmax temperature of T=0.1) over ten runs per position and compared the results with the exhaustive mutational analysis obtained from DDgPredictor (**Fig. S8**). For 5FV2, we observed that AbMPNN predictions often stick to the wild-type amino acid and other substitutions are mostly favorable according to DDgPredictor. For 3P9W, AbMPNN frequently predicts wild-type amino acids while its alternative amino acid substitutions are often unfavorable (according to DDgPredictor) and tend to be limited to a small set of choices. For instance, Tyrosine is predicted at 6 of the 13 positions. At the given Softmax temperature, AbMPNN tends to have less variability, resulting in a smaller exploration of the mutational landscape, and may miss potentially beneficial mutations. We used predictions from DDgPredictor to guide the mutations introduced by AbMPNN, either preserving the mutation or reverting to the wild-type amino acid. This combination enabled us to achieve energy reductions as low as -4.5 *kcal/mol* (**Fig. S9**).

All mutants exhibit a *T*_*m*_ within a range of 3.7°C and 5.3°C, while the wild-type *T*_*m*_ are 72.49°C and 71.54°C for VHH in complexes 3P9W and 5FV2, respectively (**Fig. 5AB**). We observed that specific CDR3 positions favor higher *T*_*m*_ (3rd for 5FV2 and 4th for 3P9W) while others are very sensitive to mutations and tend to reduce it (2nd for 5FV2 and 5th and 11th for 3P9W).

**Figure 5:**
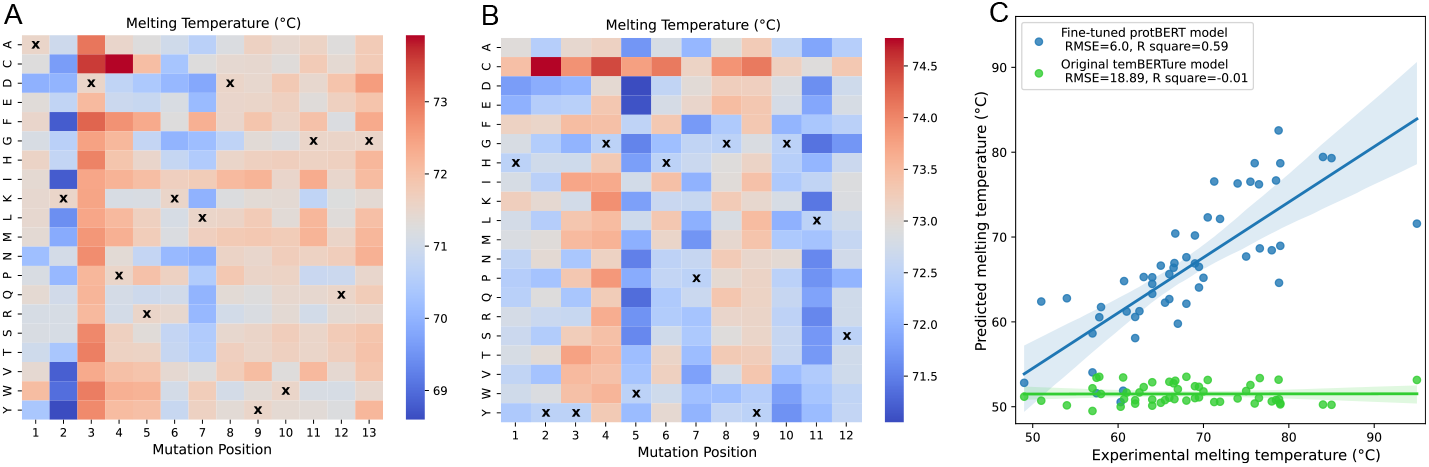
**(A-B)** *T*_*m*_ prediction for exhaustive point mutation performed on CDR3 of VHH in 5FV2 (**A**) and 3P9W (**B**). Crosses indicate the wild-type amino acid. Higher temperatures mean higher thermostability. **(C)** Comparison between the melting temperature (*T*_*m*_) determined experimentally and the one predicted by the temBERTure and fine-tuned models. RMSE=Root Mean Square Error.

Our fine-tuned model for predicting melting temperature outperforms the baseline standard model, reaching lower Root Mean Square Error (RMSE) (6.0°C vs 18.89°C), higher R-square (0.59 vs -0.01), and improved Pearson correlation coefficient (0.772 vs 0.005) (**Fig. 5C**). The predicted vs experimental curve shows a closer alignment with the ground truth, highlighting the fine-tuned model’s ability to capture the effects of mutations on *T*_*m*_ more effectively than the standard model. See **Fig. S2** for model improvement across different epochs, and **Fig. S3-S5** for the ablation study to find the best architecture and hyperparameters.

## 4 Discussion

In this work, we developed a deep-learning framework for VHH CDR3 design via affinity maturation, modeling how mutations affect backbone conformation, VEGF binding affinity, and VHH thermostability. To enhance the reliability of our ΔΔ*G*_*Bind*_ predictions, future improvements will incorporate molecular dynamics simulations to better capture protein complex conformational variability. Moreover, the accuracy of mutant protein structure models remains an active area of research. Studies suggest AlphaFold2 [12] is not reliable for predicting mutation-induced changes in protein stability or function, as it can model folded structures for mutations that experimentally yield unfolded proteins [28, 20]. While our VHH *T*_*m*_ model outperforms the state-of-the-art, its sensitivity must be improved to fully capture the effects of single-point mutations on VHH stability. Finally, the model was trained exclusively on folded VHHs due to the lack of experimentally measured *T*_*m*_ for non-folding variants. Availability: The source code and model weights are available from the corresponding author upon request.

## Acknowledgements

This work was supported and funded by National Institute for Research in Digital Science and Technology (Inria) at Inria Paris Center.

## Supplementary Information

**Figure S1:**
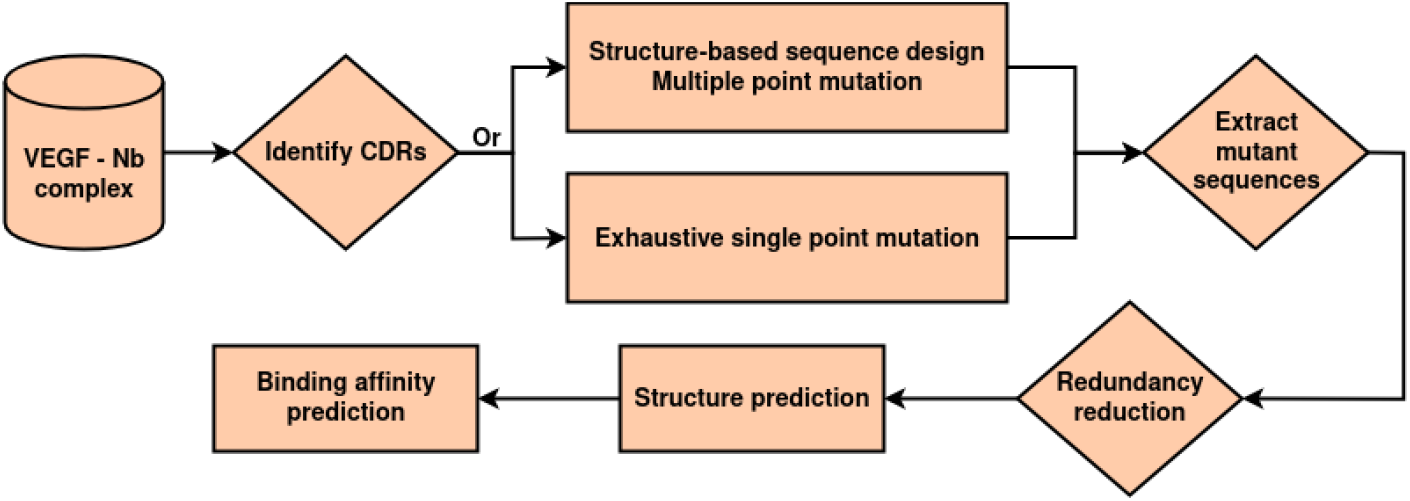
Wild-type based maturation pipeline. Multiple point mutations or exhaustive single point mutations are performed on the CDR3. After redundancy reduction, the structures are generated from the mutant sequences, and the binding affinity changes are predicted.

**Figure S2:**
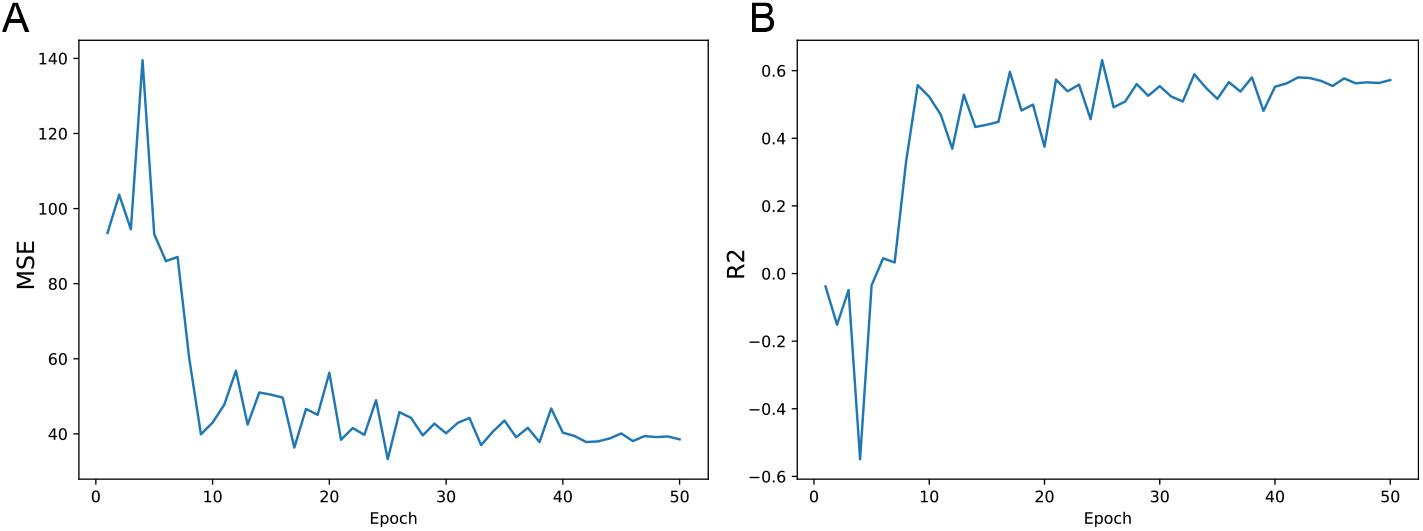
Evolution of **(A)** the Mean Square Error (MSE) and **(B)** *R*^2^ over finetuning protBERT. The X-axis is the epochs and the Y-axis is the MSE or *R*^2^.

**Figure S3:**
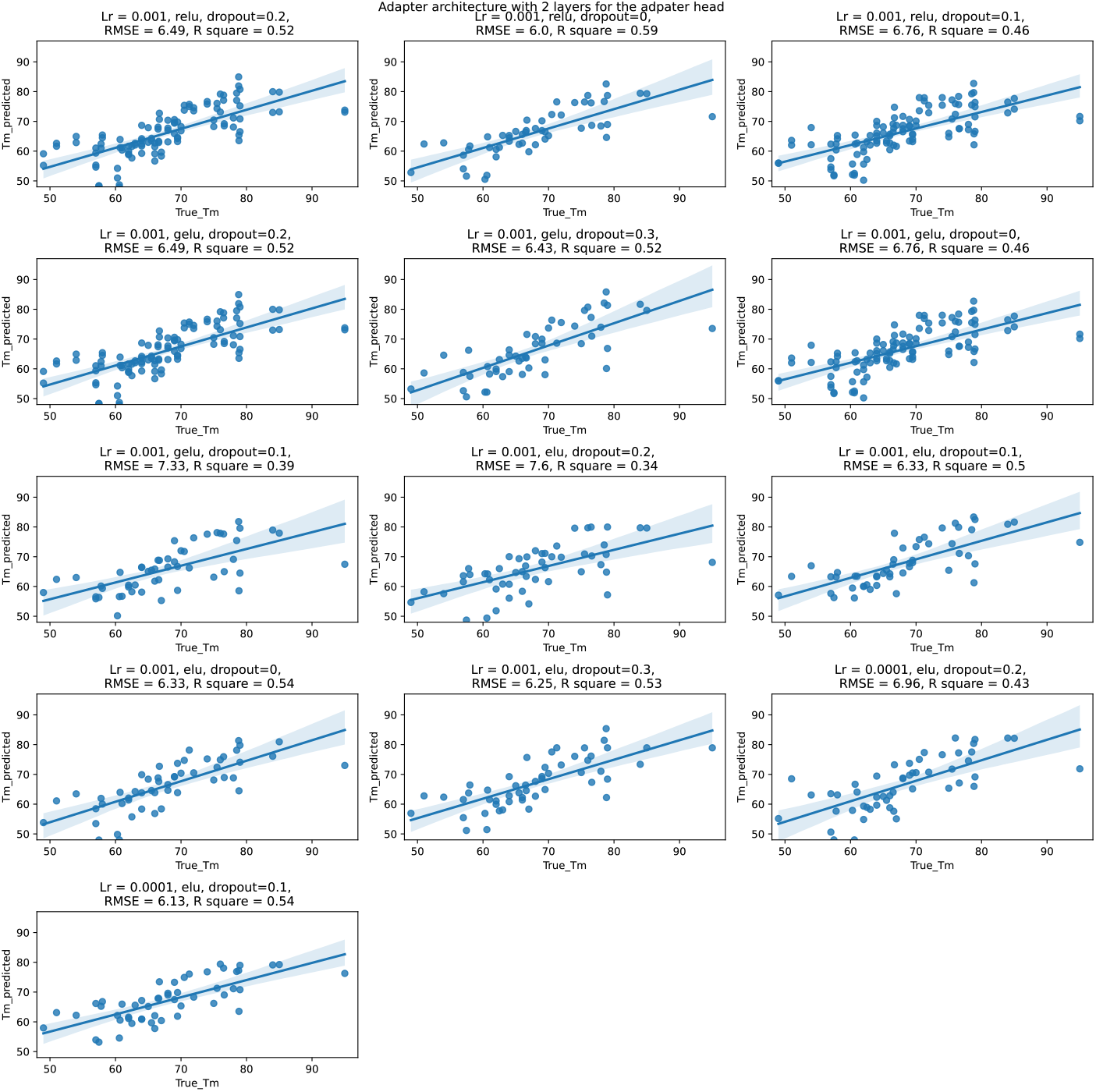
The performance of VHH melting temperature predictor on the test set. The finetuning of the parameters was performed on an adapter architecture on protBERT and with 2 layers in the adapter head. The learning rate (Lr), dropout, and activation function in the adapter heads were changed. RMSE stands for root mean square error.

**Figure S4:**
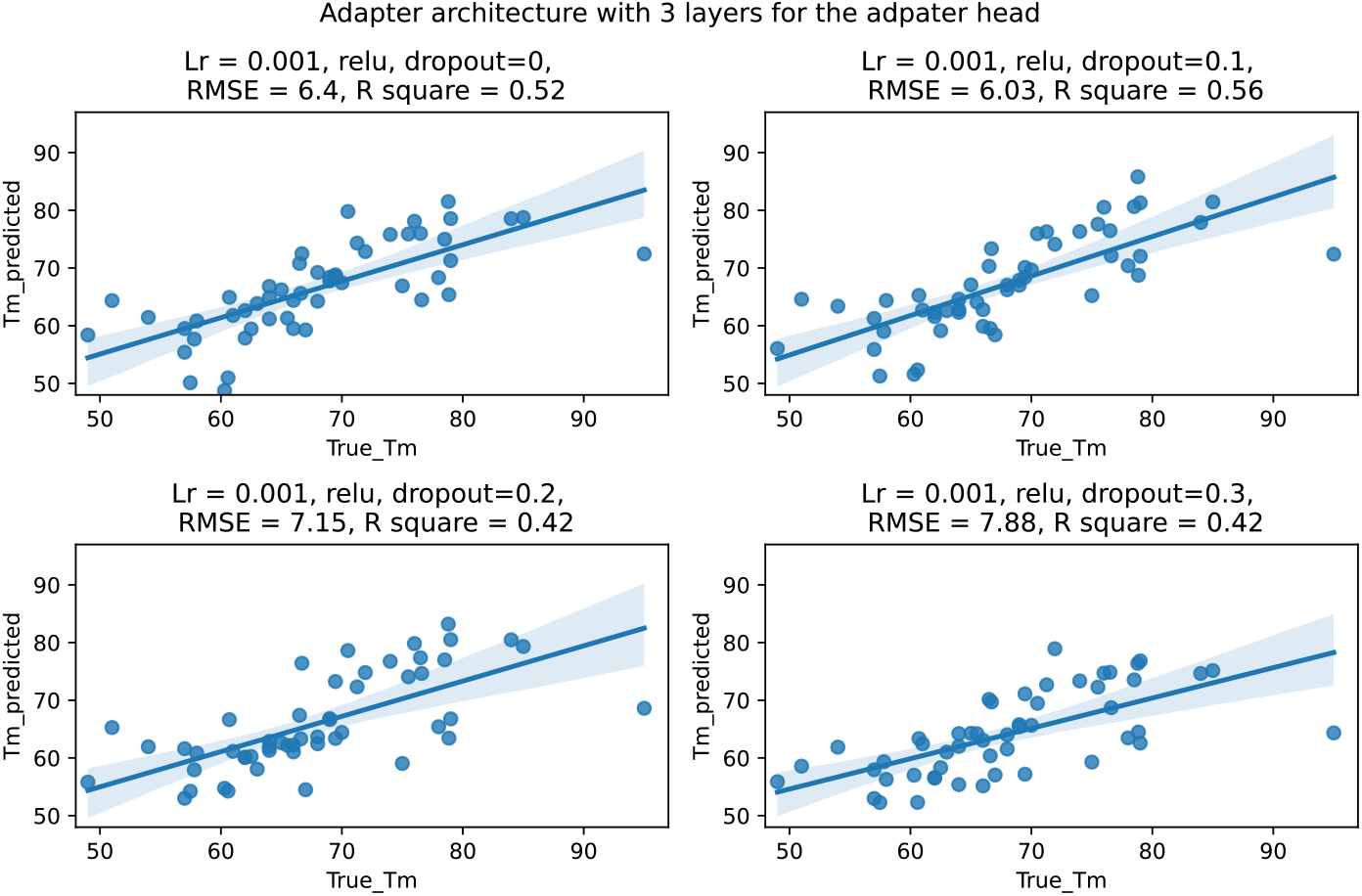
The performance of VHH melting temperature predictor on the test set. The finetuning of the parameters was performed on an adapter architecture on protBERT with 3 layers in the adapter head and a learning rate of 0.001. The dropout was changed. RMSE stands for root mean square error.

**Figure S5:**
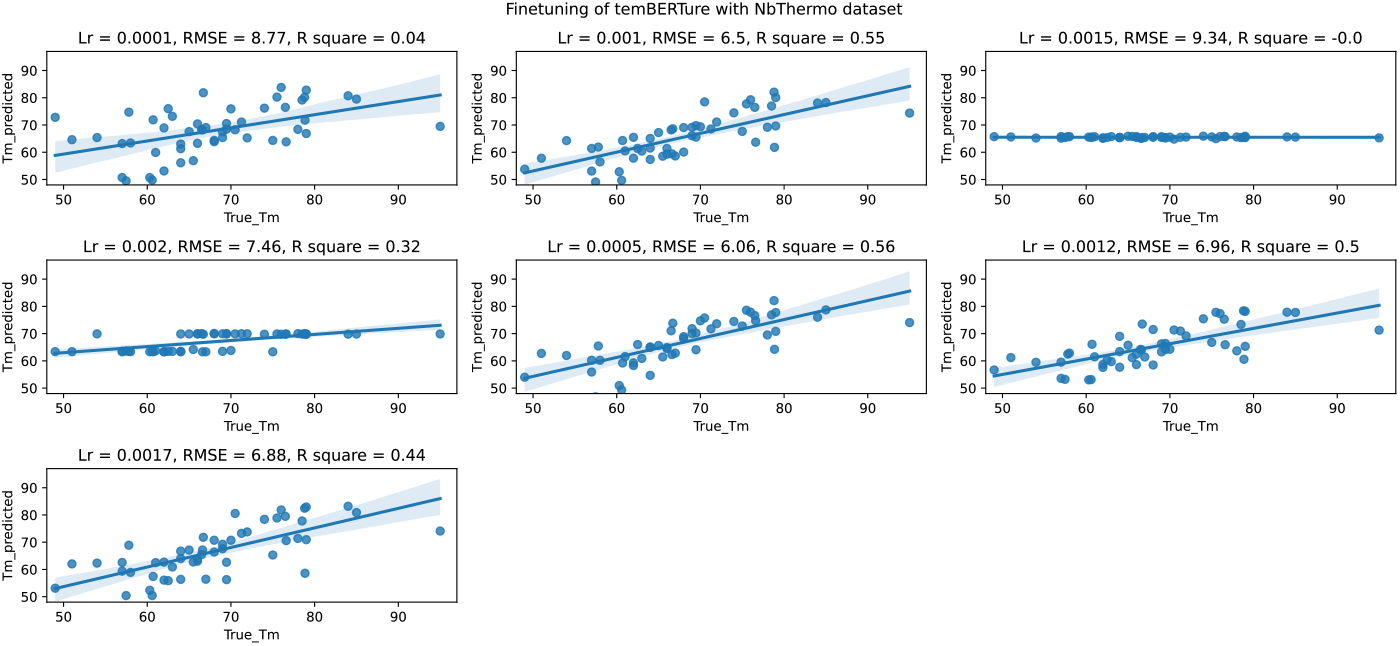
The performance of VHH melting temperature predictor on the test set. The parameters were finetuned on the temBERTure best model (replica 2). The learning rate (lr) was changed. RMSE stands for root mean square error.

**Figure S6:**
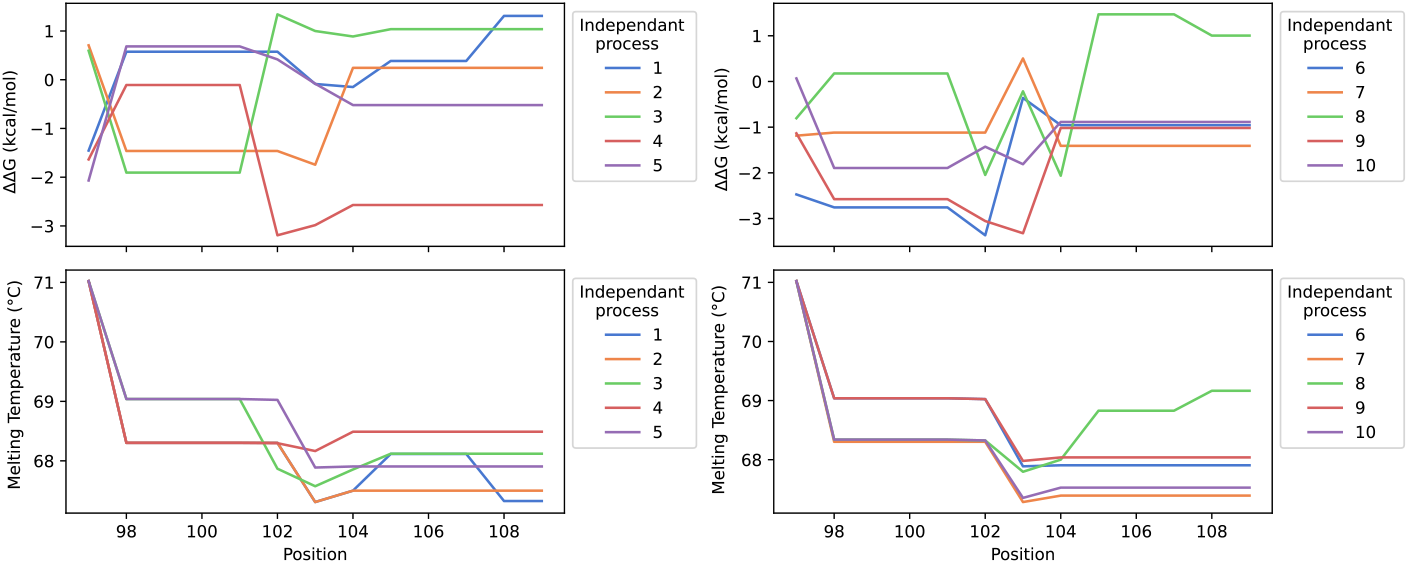
Evolution of the predicted changes in binding affinity (top) and melting temperature (bottom) through mutant positions within the CDR3 of the 5FV2 complex. Each curve corresponds to an independent process. The X-axis position corresponds to the amino acid number with respect to the first amino acid of the nanobody. CDR3 region has been identified to be between amino acids 97 and 109.

**Figure S7:**
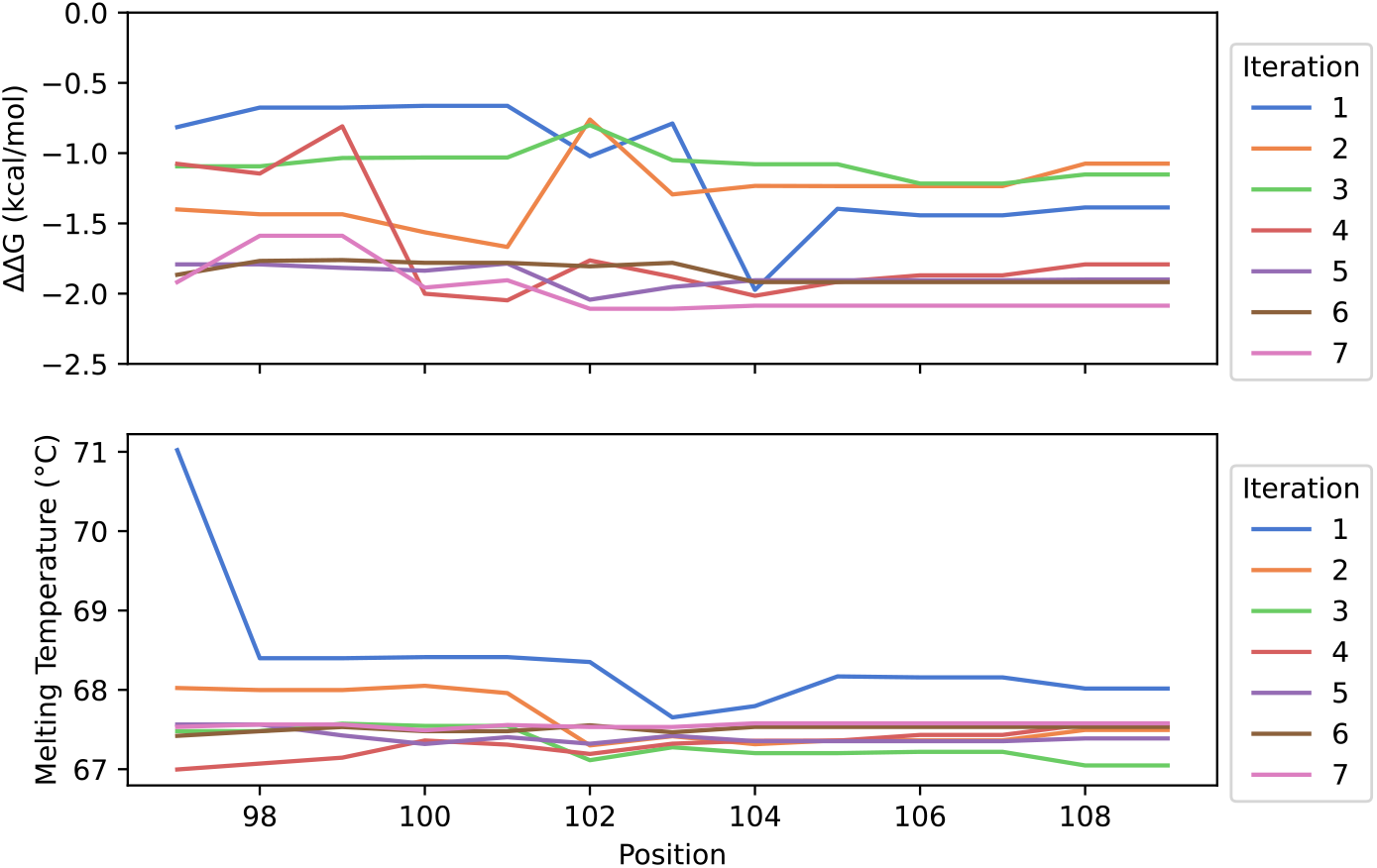
Evolution of the mean of predicted changes in binding affinity (top) and the mean of predicted melting temperatures (bottom) through mutant positions within the CDR3 of the 5FV2 complex for 7 iterations. The mean is calculated over the 10 independent runs. The mean of changes in binding affinity (top) reaches a lower value at the end of the last iteration. The mean melting temperature (bottom) remains in a 2°C range for all iterations except the first one, which reaches slightly higher values.

**Figure S8:**
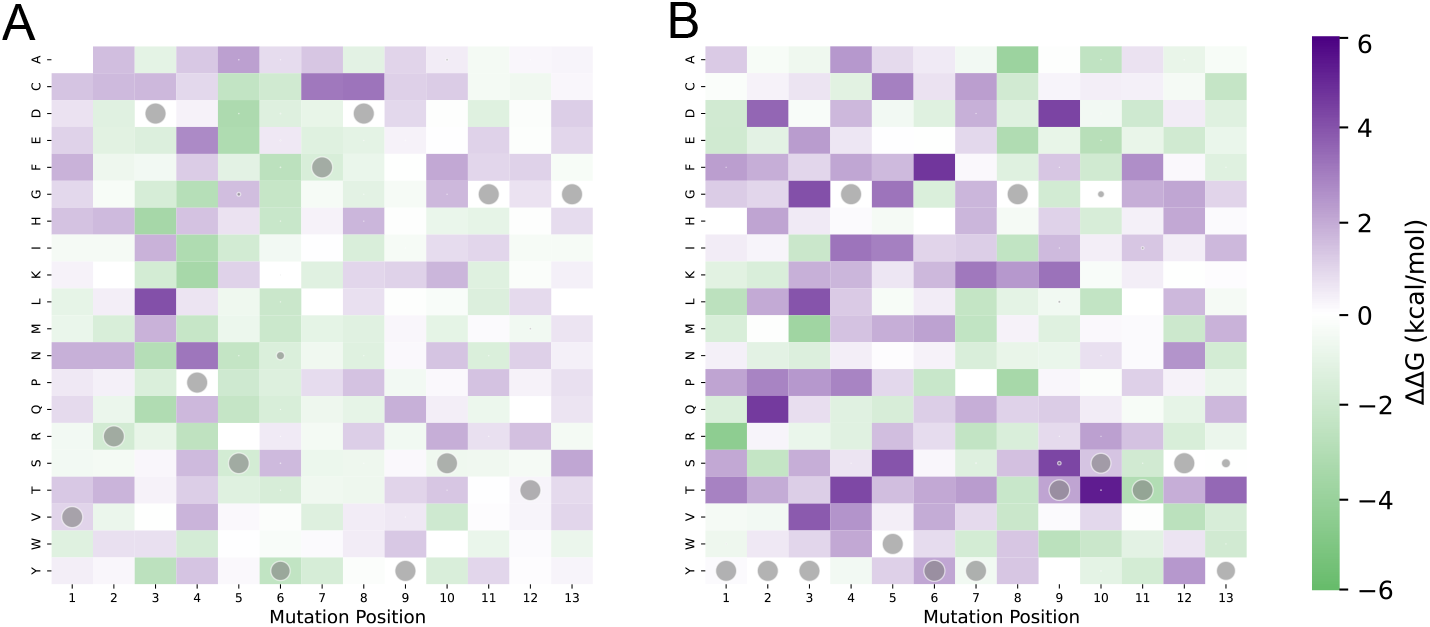
Predicted ΔΔ*G*_*Bind*_ for exhaustive single point mutation on the CDR3 of VHH in 5FV2 **(A)** and in 3P9W **(B)**. Gray circles represent the mean probability of amino acid substitution over 10 independent runs by AbMPNN for each amino acid position.

**Figure S9:**
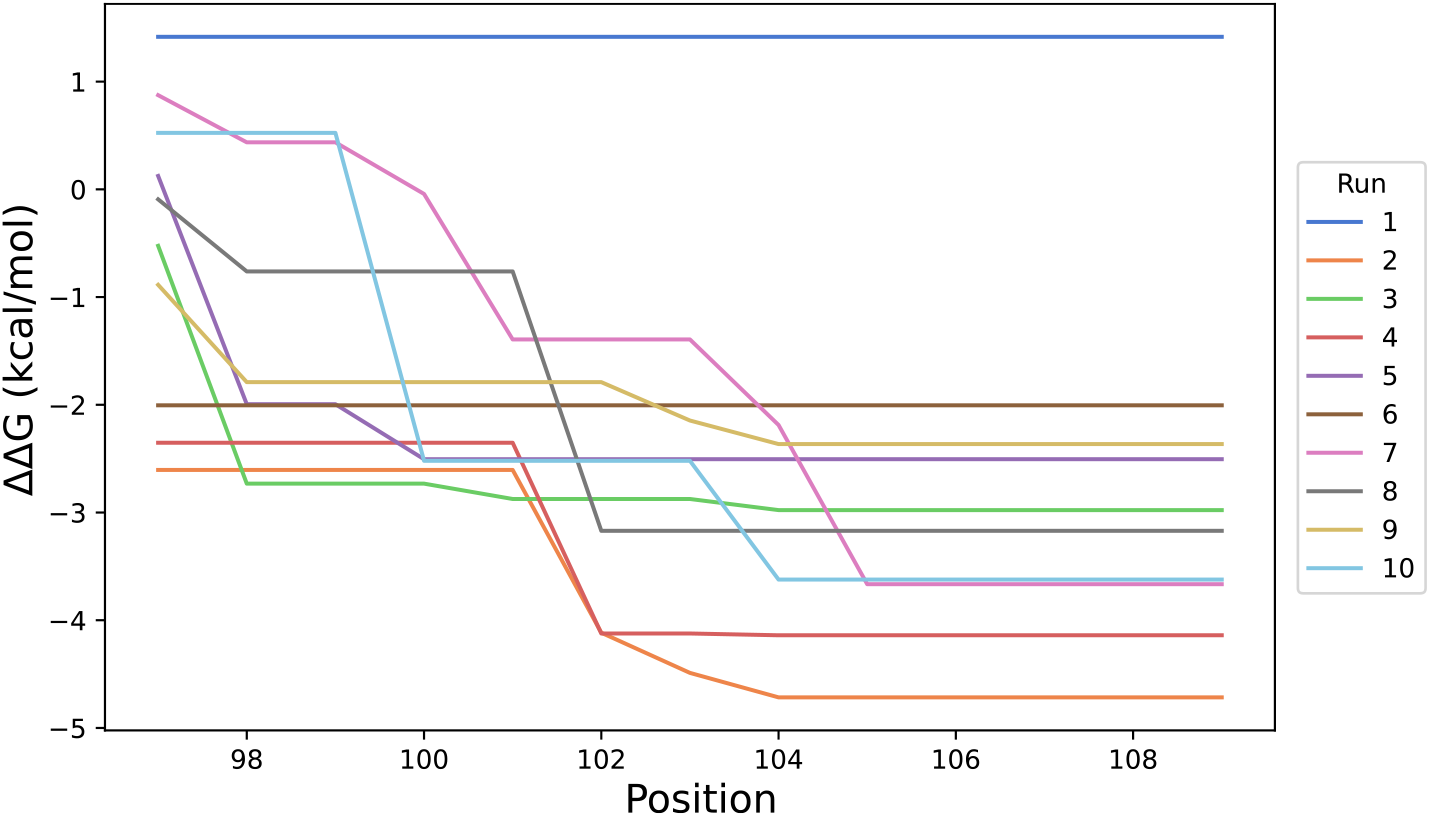
Evolution of the predicted ΔΔ*G*_*Bind*_ through CDR3 mutant positions of VHH in 5FV2 complex for the guided mutation process over 10 independent runs. AbMPNN was used instead of ProteinMPNN. The affinity maturation stops in the first amino acid position for processes 1 and 6, while for all the other processes, more favorable mutants are reached at later positions. The X-axis is the position corresponding to the amino acid number with respect to the first amino acid of VHH. CDR3 region has been identified to be between amino acids 97 and 109.

